# The trophic pyramid revisited: most animal classes have more predator than herbivore species

**DOI:** 10.1101/2022.09.14.507913

**Authors:** Tamir Klein, Stav Livne-Luzon, Uri Moran

**Affiliations:** Department of Plant & Environmental Sciences, Weizmann Institute of Science, Rehovot, Israel

**Keywords:** Multi-trophic interactions, vertebrates, invertebrates, diet, food source, mammals, insects, predators

## Abstract

According to the trophic pyramid, a large array of herbivores (primary consumers) feed an increasingly narrower array of predators, from secondary consumers to top predators. However, the partitioning between herbivory and predation across the animal kingdom has not yet been tested at the global scale.

Here, we use a survey of 33,762 animal species across ten major taxonomic groups (five vertebrates and five invertebrates) to partition food sources at the class and phylum levels. We use this partitioning, together with class-level biomass estimates, to create a global trophic pyramid of biomass.

We show that: (1) the diet of eight of the ten groups of animals is dominated by prey rather than plants, accounting for 64-99% of the diet mass; (2) collectively across the terrestrial and marine environments, secondary consumers (invertivores; ∼1200 Mt C) have higher biomass than primary consumers (herbivores; ∼500 Mt C); (3) the two major exceptions, feeding mostly on plants, are mammals and insects; the latter form the major food source for terrestrial animals.

For animal species in most classes, plants are not a food source, but rather invertebrates, mostly arthropods. Secondary consumers (invertivores) link primary consumers and top predators, and are hence pivotal to almost all food-webs.

## Introduction

Since its inception nearly a century ago, the trophic pyramid has been one of the cornerstones of ecology (Elton 1927, Bodenheimer 1938, Lindeman 1942). The pyramid of numbers claimed that the abundance of animals in the base of food chains is larger than the abundance of animals at the end of such chains (Elton 1927). The concept was then expanded to the pyramid of biomass, whereby a large biomass of producers (plants) is feeding a smaller biomass of herbivores, which in turn feed a yet smaller biomass of carnivores, and so on up to top predators (Bodenheimer 1938). Applying the rules of thermodynamics, it became clear that each trophic level of the pyramid should retain only a fraction of the energy of the level below, giving rise to the pyramid of energy (Lindeman 1942). Further, it was suggested that the slope of the pyramid, defined as the increase in biomass between successively lower trophic levels, may reflect ecosystem stability (Raffaelli 2002). Evidence contrasting the bottom-heavy shape of trophic pyramids came from a survey of plankton types in the North Atlantic, suggesting an ‘inverted pyramid’ (Buck et al. 1996). Similar observations were found in a few other ecosystems (Fath and Killian 2007). On the other hand, another study claimed that these contrasting evidence indicate overestimation of predator abundance or energy subsidies (Trebilco et al. 2013). A meta-analysis of 72 food webs concluded that inverted pyramids, of species richness in that case, should be rare (Turney and Buddle 2016).

Recently, it was argued that inverted, top-heavy, pyramids of biomass can be stable and prevalent in the natural world (McCauley et al. 2018). Such pyramids can emerge from a variety of mechanisms, either endogenous or exogenous to the ecosystem. The first such mechanism relates to transfer efficiency across trophic boundaries (e.g., biomass transfer >10%, which is typically assumed; Pauly and Christensen 1995), which can be increased through increases in the edibility and nutrient content of foods or higher foraging efficiency of consumers (McCauley et al. 2018). A second major mechanism is rapid turnover rates, e.g., whereby herbivores reproduce faster than their predators. Other endogenous mechanisms include higher predator access to different prey pools, facilitated by increased resource production; and omnivory by predators (McCann and Hastings 1997) as means to feed on both herbivores and producers, thereby supporting increase in their biomass. Exogenous mechanisms of inverted pyramids include the vectoring of distant subsidies to predators and consumers, or the sourcing of food outside community boundaries in favor of local predators. For example, top-heavy coral reef communities were described in which mobile consumers draw energy from neighboring pelagic communities (McCauley et al. 2018). Subsidies operate in both space and time, and hence historical subsidies also exist, whereby high predator biomass can be facilitated by herbivores that were previously present in the food web. Inverted pyramids were identified in specific marine and estuary environments (Buck et al. 1996, Fath and Killian 2007), as well as in freshwater ecosystems including streams, lakes, and ponds (Fath and Killian 2007). A few examples were described in terrestrial ecosystems, yet at far lower incidence (Fath and Killian 2007, McCauley et al. 2018). However, the validity of the fundamental concept of the trophic pyramid has never been tested across diverse terrestrial and marine animal classes simultaneously, and particularly not on a global scale.

Here we compiled a dataset of diets of 33,726 animal species across ten major taxonomic groups: the five vertebrate classes, and the five major invertebrate groups. Based on the available literature, we hypothesized that the classic, bottom-heavy trophic pyramid is true for the terrestrial environment, whereas the inverted, top-heavy pyramid is true for the marine environment. Consequently, the shape of the global trophic pyramid should reflect the biomass ratio between them.

## Materials and Methods

### Data acquisition and dataset construction

We compiled a dataset of diets of 33,726 animal species across ten major taxonomic groups: the vertebrate classes mammalia, aves, reptililia, amphibia, chondrichthyes and osteichthyes (jawed fish and bony fish, respectively, grouped here under fish), and the invertebrate classes arachnida and insecta; the subphylum crustacea; and the phyla mollusca and cnidaria. We selected these groups since they represent the vast majority of animal diversity: the five vertebrate groups collectively represent 96% of all vertebrate species and the five invertebrate groups collectively represent 92% of all invertebrate species (Chapman 2009, Ruggiero et al. 2015). In each group, we obtained information either for almost all species (for mammals and birds; Wilman et al. 2014), or for the largest array of species available in the literature, with a minimum of ∼500 species per group. In most groups, we ensured representation for all orders and most families. For each species, we used its full diet partitioning by mass, rather than a single-term ‘herbivore’, ‘carnivore’, etc. For example, for the common blackbird, *Turdus merula*: 20% grains, 20% fruits, 50% invertebrates, and 10% vertebrates (Wilman et al. 2014). Such species-level data were not used directly but were rather used to create a class-level partitioning of food sources (Table 1). Doing so also enabled us to avoid the term ‘omnivore’, which encapsulates many diet sources, all of which are important for our analysis (Pineda-Munoz and Alroy 2014). We note, however, that our analysis regards the partitioning of diet at the class or phylum level, and not at the species level, where mixing of diet types is also important (Pineda-Munoz and Alroy 2014). Next, we multiplied the class-level partitioning by a class-level biomass estimate, to produce a biomass estimate for the role of each food source in feeding each of the ten groups. While being simplistic, our class-level approach allowed us to circumvent the need for species-specific data on average body mass and population size (Fig. S1).

**Table 1.** An example of our calculation approach. We used species-specific food source partitioning, demonstrated below for seven bird species (rows), to calculate the relative contribution of each of eight food categories (columns) at the class level. In other words, the class-level ratios are those for a theoretical “average bird”, based on 9993 bird species.

| Species | Plant | Plant<br>derived | Fungi | Detritus | Parasite<br>invertebrate | Animal<br>invertebrate | Parasite<br>vertebrate | Animal<br>vertebrate |
| --- | --- | --- | --- | --- | --- | --- | --- | --- |
| Abeillia<br>abeillei | 90 | 0 | 0 | 0 | 0 | 10 | 0 | 0 |
| Babax<br>koslowi | 60 | 0 | 0 | 0 | 0 | 40 | 0 | 0 |
| Cacatua<br>alba | 90 | 0 | 0 | 0 | 0 | 10 | 0 | 0 |
| Dacelo<br>tyro | 0 | 0 | 0 | 0 | 0 | 100 | 0 | 0 |
| Egretta<br>sacra | 0 | 0 | 0 | 0 | 0 | 50 | 0 | 50 |
| Falco<br>alopex | 0 | 0 | 0 | 0 | 0 | 30 | 0 | 70 |
| Galbula<br>dea | 0 | 0 | 0 | 0 | 0 | 100 | 0 | 0 |
| ... up to<br>$n = 9993$ | | | | | | | | |
| **Class-**<br>**level**<br>**ratios** | **36** | **0** | **0** | **0** | **0** | **56** | **0** | **8** |

Currently, body mass and abundance (population size) data are available only for bird species. We used these data to produce a species-weighted diet source biomass partitioning, for comparison with our aforementioned simplistic approach. We merged the bird species’ diet and body mass dataset (Wilman et al. 2014) with the new bird species’ abundance dataset (Callaghan et al. 2021). The merged dataset included 7,548 bird species, i.e., ∼75% of all known bird species (Table S1). For each species, we multiplied body mass with abundance, to produce the total species biomass. We then partitioned the total species biomass among its respective diet sources. Next, we summed the biomass of all birds for each diet type and calculated its share among the total global bird biomass. Finally, we compared the results of this analysis to that of our general approach for the ten animal groups.

Data sources for animal species diets included: (1) Comprehensive data for most extant mammals (5397 species) and birds (9993 species), from Wilman et. al. 2014; (2) data for >8,000 species of fish from FishBase (see below); (3) data for ∼3,000 species of lizards for which dietary data is available, from Meiri 2018; (4) data from various, confirmed, sources reported on Wikipedia; (5) the Encyclopedia of fauna and flora in the Land of Israel; and (6) peer-reviewed papers. We gathered dietary data for no less than 480 (483-9993) species in each of the following animal groups: mammals, birds, reptiles, amphibians, fish, insects, arachnida, crustacea, mollusca, and cnidaria. In the case of the large class of insects, we strategically focused on the six largest orders, collectively containing 93% of all known insect species. Still, our sampling rate (by percentage) was lower for insects than for other groups. Nevertheless, it was sufficient to be representative of insects’ food sources (Fig. S2; See under *Statistical analysis*). We collected dietary data for no less than ∼250 (252-1404) insect species in each of the following orders: Coleoptera, Diptera, Hemiptera, Hymenoptera, Lepidoptera, and Orthoptera. Notable omissions in our dataset are the worm phyla annelida and nematoda, collectively representing 3% of all invertebrate species. Although the worm phyla contain many important detritivore species, information on their phylogeny and diet sources is still incomplete.

Diet data classification: Where available, we classified dietary data according to the following 17 categories, adapted from Wilman et al. (2014) (‘Vert’ is short for ‘Vertebrates’, ‘Invert’ is short for ‘Invertebrates’): ‘Vert (general)’, ‘Vert (mammals/birds)’, ‘Vert (reptiles/amphibians)’, ‘Vert (fish)’, ‘Vert (scavenger)’, ‘Vert (parasite)’, ‘Invert (general)’, ‘Invert (scavenger)’, ‘Invert (parasite)’, ‘Plants (general)’, ‘Frugivore’, ‘Nectarivore’, ‘Granivore’, ‘Fungivore’, ‘Detritivore’, ‘Plant derived’ (the latter category is applicable in cnidaria only), and ‘Other’. For the purpose of producing a comparative analysis across ten animal groups, we finally narrowed down to 8 diet classes: vertebrate (of all forms, including scavenging, excluding parasite), parasite on vertebrate, invertebrate (of all forms, including scavenging, excluding parasite), parasite on invertebrate, fungi, detritus, plant (any plant part including fruit, nectar, and seed), and plant-derived (in the case of cnidaria, where many species (e.g. of corals) obtain their resources directly from symbiotic algae). Whenever available, we specified whether the data was for adult or juvenile, and for male or female. As a rule, we tried to have data as diverse as possible for each animal group. To maintain this diversity, for the most part we did not include more than 16 species from a single genus. An additional group, ‘Insects combined’, was created by collating data for male adults from the six insect orders. In Lepidoptera, data was often available only for juveniles, while adults in this order feed on nectar, pollen, or even lack mouth parts and do not feed at all (Altermatt and Pearse 2011). Another noteworthy point is that among 647 crustaceans, 35 of the selected species were from the highly important order Euphasiacea (krill) and 117 of selected species were from the Copepoda orders Siphonostomatoida, Cyclopoida, and Harpacticoida. We found that while krill feeds mostly on phytoplankton, many Copepoda species are parasitic, mostly on fish.

For each species, the data source reference was specified, and, where available, the description of the diet (dietary comments) was added. The dietary comments were copy/pasted (verbatim) from the data source. The dietary value assignment for each species was made according to the comments’ section, giving a total of 100%. Fish dietary data was extracted from FishBase, using their most general classification, ‘Food I’: ‘detritus’, ‘plants’, ‘zoobenthos’, ‘zooplankton’, ‘nekton’, and ‘other’. We assumed nekton to be comprised of 80% fish and 20% mollusca. As can be seen on the FishBase food items table, zoobenthos includes only invertebrates. The ‘other’ category, which was negligible (18 out of 8367 species), was ignored, by subtracting it from the sum of species. Generally, ‘Plankton’ (when unspecified) was assigned to a ratio of 5:1, zooplankton: phytoplankton, as advised in Bar-On et al. (2018). In many other cases, explicit zooplankton: phytoplankton ratios were specified. In lichen, as advised by Kristin Palmqvist, the ratio between algae and fungus was taken to be 1:9 in foliose lichen, and 1:1 in gelatinous lichen. When lichen type wasn’t specified, we took it to be gelatinous.

### Location-specific animal diet partitioning

Once data for species occurring globally from the above-mentioned animal groups was collected, we set out to compare the data to species occurring in natural reference ecosystems, those of the Kruger National Park (KNP) in South Africa, and the Galapagos islands (Ecuador). The two regions share a long history of conservation and are hence among the least affected by humans (Plumptre et al. 2021). These unpublished species lists were created during the past two decades by local expert scientists, primarily G. Zambatis in KNP, and A. Chiriboga, D. Ruiz, N. Tirado-Sanchez, and S. Banks in the Galapagos. Species’ dietary lists for KNP and the Galapagos were created by extracting data for species that appeared both in these two locales and in the global dietary lists. Data for species that appeared in the KNP and Galapagos lists, but did not appear in the global lists, was imputed from the global lists, when possible, in the following way: when data was not available for a KNP or Galapagos species, we looked for data for species from the same genus in the global lists. In case there were data in the global lists for at least 4 species from the same genus, and all of these species had identical diets, data were imputed from the global lists into the KNP or Galapagos species lists for that particular species. We are aware that in the case of terrestrial animals in the Galapagos, this assertion is limited due to adaptive radiation during island colonization, and in any case, only a small minority of species were added to the analysis this way. Only animal groups that had dietary data for ten species or more in KNP and Galapagos were considered in our analysis.

### Additional datasets

To test if herbivory is related to higher body mass and lower metabolic rate, we collected data on food sources of dinosaurs. Data for dietary habits of 102 non-Avian dinosaur species (termed here “dinosaurs” for simplicity) was collected from various sources reported on Wikipedia. This dataset included 38 species of the order Ornithischia (clades Ankylopollexia, Ornithopoda, and Hadrosauromorpha) and 64 species of the clade Saurischia, which had food source estimate based on dental morphology, and were classified into the 17 categories mentioned above. An effort was made to collect data from as many families as possible.

Human diet source partitioning was calculated using the National Geographic Society application “What the world eats” (https://www.nationalgeographic.com/what-the-world-eats/), which is based on FAO data. Partitioning was made on daily amounts of the global mean human consumption of different food categories (g d^−1^) calculated for 2011. Seafood was divided between fish (animal vertebrate, 80%) and crustacea and mollusca (animal invertebrate, 20%). Milk and eggs were categorized as parasites on vertebrates.

### Partitioning of animal diet by biomass and construction of global food webs

Estimates for the total biomass of animals from specific groups were taken from Bar-On et al. (2018). Marine arthropods were considered as crustacea. One exception was the total biomass estimate for amphibians, where the authors admitted that their approach produced an over-estimation (Bar-On et al. 2018). Instead, we made a comparison with another ectotherm class, i.e. reptilia, which had a higher-certainty estimate. While being similar in the number of species, individual mean body mass is one order of magnitude lower in amphibians than reptiles (1 vs. 10 g C; Feldman et al. 2016), whereas population density is roughly an order of magnitude higher. Together, this meant an equal class-level biomass estimate across amphibians and reptiles. The group-specific total biomass value was then divided among diet categories, to yield estimates of total biomass of herbivores, invertivores, vertivores, and so forth, for each group. While these estimates are very useful for our trophic level analysis, they have two inherent problems: (1) they preclude omnivores, or essentially any type of mixed diet at the species level, and (2) they assume a homogeneous body mass across trophic levels of the same class, e.g. in the case of the herbivore hare, the invertivore hedgehog, and the vertivore wild cat, which are roughly of similar biomass (0.3-3 kg C). Considering species with mixed dietary types (problem 1), while they are not strictly represented in our trophic hierarchy (as also in the classic trophic pyramid; Odum et al. 1968), by dividing their biomass across the trophic levels, we do account for their multiple trophic roles (Table 1). Considering the assumption of group-specific, homogeneous, body mass (problem 2), this is not to be mistaken with the well-studied predator-prey mass ratio (Trebilco et al 2013), which can span over multiple orders of magnitude. Rather, it relates to the variation of body mass across dietary types within a group, which is far smaller. For example, based on Wilman et al. (2014), 50% of all bird species have a body mass between 15 and 130 g, and 80% of all bird species are between 9 and 530 g. Variation is larger in mammals, with 50% of species between 20 and 580 g, and herbivores being typically bigger than carnivores. Still, within-group body mass variation can be large: e.g., many small mammals (<1 kg) are either insectivores or granivores, and many larger mammals (>30 kg) are either carnivores or herbivores (Pineda-Munoz et al. 2016). Even when considering the predator-prey mass ratios (including across classes), they vary widely but concentrate close to 1.0 (Brose et al. 2006).

Across many food webs, a tradeoff exists between body size and population size (Barnes et al. 2010). As a result, variations in the contribution of each species to the total biomass of its class are reduced. Predator-prey mass ratios are high in freshwater ecosystems, yet much lower in vertebrates in marine and terrestrial ecosystems and more so among invertebrates, which make up most of the biomass globally (Brose et al. 2006). The partitioning of diets across the animal kingdom, expressed also as biomass, facilitated the construction of global food webs, for Earth’s marine environment and terrestrial environment. In each animal group, we accounted for herbivores, invertivores, vertivores, and detritus feeders, as in the hierarchy by biomass. To this, we added more detail, in the form of relationships among specific groups, e.g. the biomass of crustacea feeding on fish. Doing so yielded food webs that described the flow of biomass from producers to herbivores, on to invertivores and on to vertivores.

### Statistical analysis

In the insect class our data was estimated to cover ∼0.3% of the estimated number of described species of the class. Other groups with low coverage were arachnida, mollusca, and crustacea (0.5%, 0.9%, and 1.3%, respectively). Species coverage in cnidaria and amphibia (5.7% and 6.0%) were lower than in fish and reptilia (27.2% and 39.3%). To test the validity of our results on diet partitioning, we used the birds dataset, with almost full coverage (as in mammalia too; 98.4%), and tested how representative was such a relatively small sample. To do so, we repeatedly (10,000 times) sampled a fraction of ∼0.3% of the entire dataset we have on birds and calculated the diet partitioning in the small sample group of birds. Then, we averaged the diet partitioning in the 10,000 repeats. We ran the simulation twice: (1) including all birds’ orders. (2) including only the top 14 birds’ orders (Passeriformes, Apodiformes, Piciformes, Charadriiformes, Psittaciformes, Columbiformes, Galliformes, Accipitriformes, Strigiformes, Anseriformes, Gruiformes, Coraciiformes, Cuculiformes, Procellariiformes). The latter option simulated our strategic sampling within the six largest insect orders, collectively containing 93% of all insect species. This process was done using an independent script using the software “R” (Oksanen et al. 2017, R Core Team, 2018, version 4.0.3).

We compared the diet partitioning of the animal groups from the global dataset to that collected from the Galapagos Islands and Kruger National Park. The data on diet partitioning in each of the animal groups from each site was treated as a community-by-species matrix where the different food partitionings were treated as species from different sites (Global/ Galapagos/ Kruger). We then analyzed this data-table using non-metric multidimensional scaling (NMDS) based on Bray-Curtis dissimilarities, using the functions “metaMDS” and visualizing by “orditorp” implemented in the “Vegan” package (R Core Team, 2018, version 4.0.3).

## Results

Prey, and not plant, contributed the highest proportion of diet source within the animal kingdom (Fig. 1). In eight of ten groups, animals (prey) contributed 64-98% of the diet, and specifically, invertebrates contributed from 43% of the diet (in crustacea) to 86% (in amphibians), making invertebrate animals the single most important food source in the animal kingdom. Vertebrate animals were a smaller food source, providing up to 12% of the diet in a group, yet parasitism on vertebrates was important for arachnida (27%), and crustacea (23%). These two groups had parasites on invertebrates as well (3% each). Detritus was a small diet source for vertebrates (fish only), and a larger diet source for invertebrate groups, especially crustacea and mollusca (14% and 11%, respectively). Fungi were a very small diet source for insects, arachnida and mollusca (2%, 1% and 1%, respectively). Plants were the major diet source for insects (81%) and mammals (56%). In addition, plants were an important diet source for birds (36%) and reptiles (12%), especially turtles and some lizards. In the marine environment, algae, including phytoplankton, contributed 15-17% of the diets of fish, mollusca, and crustacea. In the case of cnidaria, many species (e.g. of corals) obtain their resources directly from symbiotic algae, and hence this diet type was termed here ‘plant derived’.

**Fig. 1.**
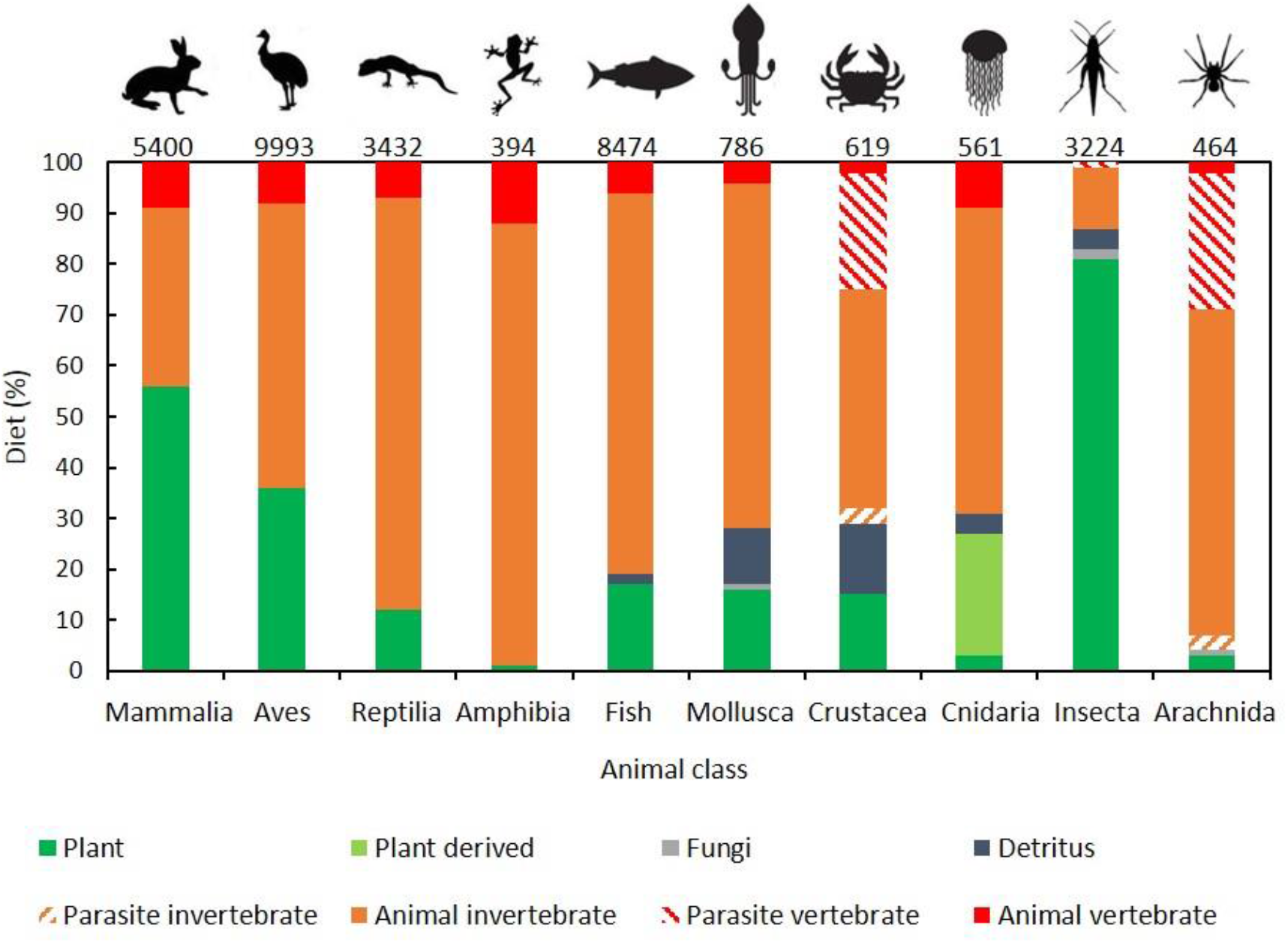
Species in most animal groups feed on prey, not plant. Partitioning of food sources among ten major animal groups. Numbers on top indicate the number of sampled species in each class/ subphylum/ phylum. Fish include all Chondrichthyes and Osteichthyes species.

To validate our observation of the primary role of prey in animal diets on a global scale, we set to test it on a local scale, in both marine and terrestrial ecosystems. Considering that current species lists in most locations on Earth miss many important species due to Anthropocene extinctions (Butchart et al. 2010, Plumptre et al. 2021), we sought locations with a long legacy of conservation. Among 5161 and 1599 animal species identified in the Galapagos Islands (Ecuador) and in Kruger National Park (South Africa), respectively, 738 and 758 species were included in our dataset. Among the Galapagos species, plants were a major food source only for reptiles, while invertebrates consisted the majority of the diet across the other groups, up to 78% (for fish), and including mammals and insects (Fig. S3). Among the Kruger savanna species, plants were a major food source for insects (77%) and birds (67%), but not for the other four groups, including mammals (43%) (Fig. S4). As shown globally, invertebrates consisted the major food source in both locations. Overall, the Galapagos and Kruger diet partitionings were similar with the global partitioning across the different animal groups, except for Kruger insects (Fig. S5). These observations are in line with several local studies on smaller numbers of species showing more carnivores than herbivores (Fath and Killian 2007). For example, carnivore: herbivore ratios in a coral reef (50 species) and in a grassland community (75 species) were 2.2 and 3.5, respectively (Fath and Killian 2007). Curiously, the global bird partitioning of diet matched the average of the partitioning patterns in Kruger and the Galapagos combined.

To construct a global division of animal biomass to trophic levels, we combined our group-specific diet partitioning with recent, improved, estimates of group-specific cumulative biomass (Bar-on et al. 2018). While it may appear simplistic, this approach allowed us to circumvent the need for species-specific data on average body mass and population size (Fig. S1). In our analysis, we assumed a homogeneous distribution of biomass across trophic levels in each group, e.g. in the case of the herbivore hare, the invertivore hedgehog, and the vertivore wild cat, which are roughly of similar biomass (0.3-3 kg C; see Methods). To allow for a global division of animals, we defined these three trophic levels, namely herbivore, invertivore, and vertivore, as representing a hierarchy similar to that of previously constructed (typically ecosystem-specific) pyramids (Schoenly 1990). To what extent is our approach valid? The only class where body mass and population size data are available is Aves, i.e., birds. A species-weighted diet source biomass partitioning for birds showed that plants, invertebrates, and vertebrates contributed 32%, 23% and 45% of the total global bird biomass (Table S1). Comparing this with the estimated partitioning (Fig. 1, second column from left) showed that: (1) the species-weighted division among herbivores and predators confirmed the estimated division, with 32% herbivory, compared with 36% herbivory in the estimate. (2) among predators, the respective contributions of invertebrates and vertebrates switched from 56% and 8% in the estimate to 23% and 45% in the species-weighted partitioning. Our trophic division took the middle-heavy shape of a violin, rather than a pyramid: invertivores, representing secondary consumers, were the largest group (1241 Mt C), larger than the herbivores (Fig. 2). We did not show the producers, which are estimated at 450,000 Mt C (Bar-on et al. 2018). Due to the dominance of marine animals (and particularly crustacea and fish; 1000 and 700 Mt C, respectively) in the global animal biomass, non-fish vertebrates were invisible in our global analysis, although they, too, formed a middle-heavy hierarchy (Fig. S6).

**Fig. 2.**
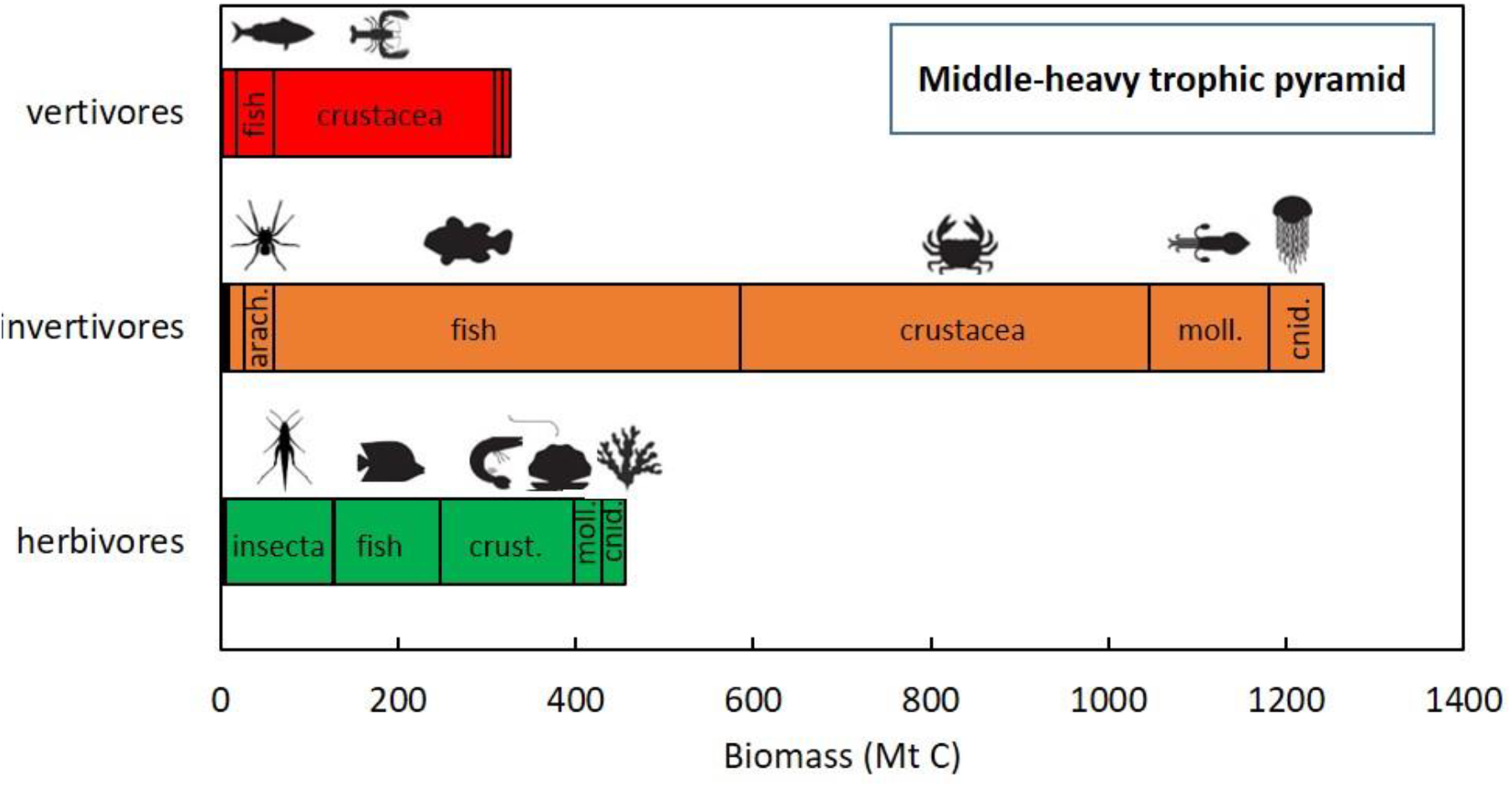
The trophic pyramid is middle-heavy: invertivores (orange; secondary consumers) are more abundant than herbivores (green; primary consumers). Estimated biomass (Megaton carbon) of herbivores, invertivores, and vertivores of each of ten major animal groups. Terrestrial vertebrates are included but relatively marginal in their biomass here, and hence are presented separately (Fig. S6) (arach., arachnida; moll., mollusca; cnid., cnidaria).

The partitioning of diets across the animal kingdom (Fig. 1), expressed also as biomass (Fig. 2), facilitated the construction of global food webs, for Earth’s marine environment (Fig. 3) and terrestrial environment (Fig. 4). Although the food web concept is inherently temporally and spatially limited and is ecosystem-specific, such a global food web, even as a conceptual exercise alone, offers insight on animal ecology at the global scale. Moreover, as many ecosystems break down and others form under the Anthropocene, a global food web can become useful. The food webs link the different groups; however, they do not present biomass flow *per se*, since the turnover of biomass is excluded here. Phytoplankton and algae feed a large biomass of crustacea and fish (150 and 119 Mt C, respectively), along with other marine invertebrates (59 Mt C). However, larger portions of these invertebrate groups (656 Mt C), and, primarily, crustacea, feed on these primary consumers (Fig. 3). Marine invertebrates, and particularly crustacea (250 Mt C) feed on fish. Yet the reverse relationship is twofold: 525 Mt C fish feed on invertebrates. The ocean’s top predators (excluding marine mammals), fish-consuming fish, are also a smaller group. Detritus feeders, which form an important component of marine food webs, sum up at 180 Mt C, are dominated by crustacea. On land, plants support a variety of herbivores, however 95% of all herbivore biomass is insects (Fig. 4). Notably, plant biomass (450,000 Mt C; not shown) is still ∼3,400-fold larger than that of all herbivores combined. Insects feed the vast majority of land animals, and primarily Arachnida and other insects. This is in line with a meta-analysis of 76 food webs from diverse biomes, showing that arachnida are the most common insectivores (Schoenly 1990). Unlike in the marine environment, detritus was not a major food source for the studied groups on land, feeding only a small part of insects, and hence excluded from Fig. 4. We note that, as in the species diet partitioning (Fig. 1), omnivory is not excluded here, although it is not explicitly presented. In practice, the biomass of any given omnivore animal species is split among its respective dietary roles.

**Fig. 3.**
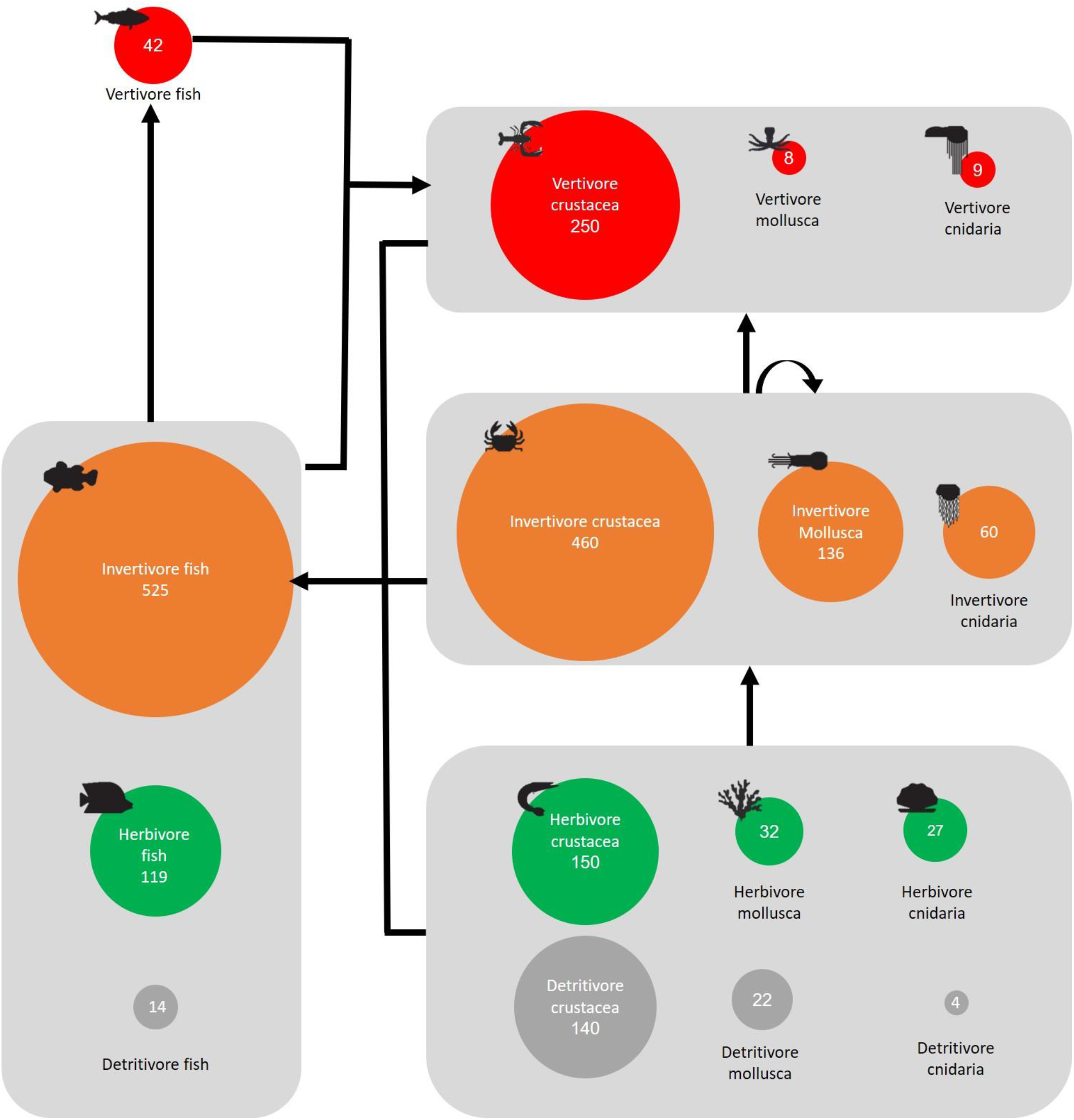
An estimated global food web for the marine environment. Numbers denote the estimated global biomass (Megaton carbon) of each group, also represented by circle area. For example, among crustacea, animals with the total biomass of 250 Mt C are vertivores, i.e. feeding on fish. In turn, all marine invertebrates (all groups within the three righthand grey areas) are potential food source for invertivore fish.

**Fig. 4.**
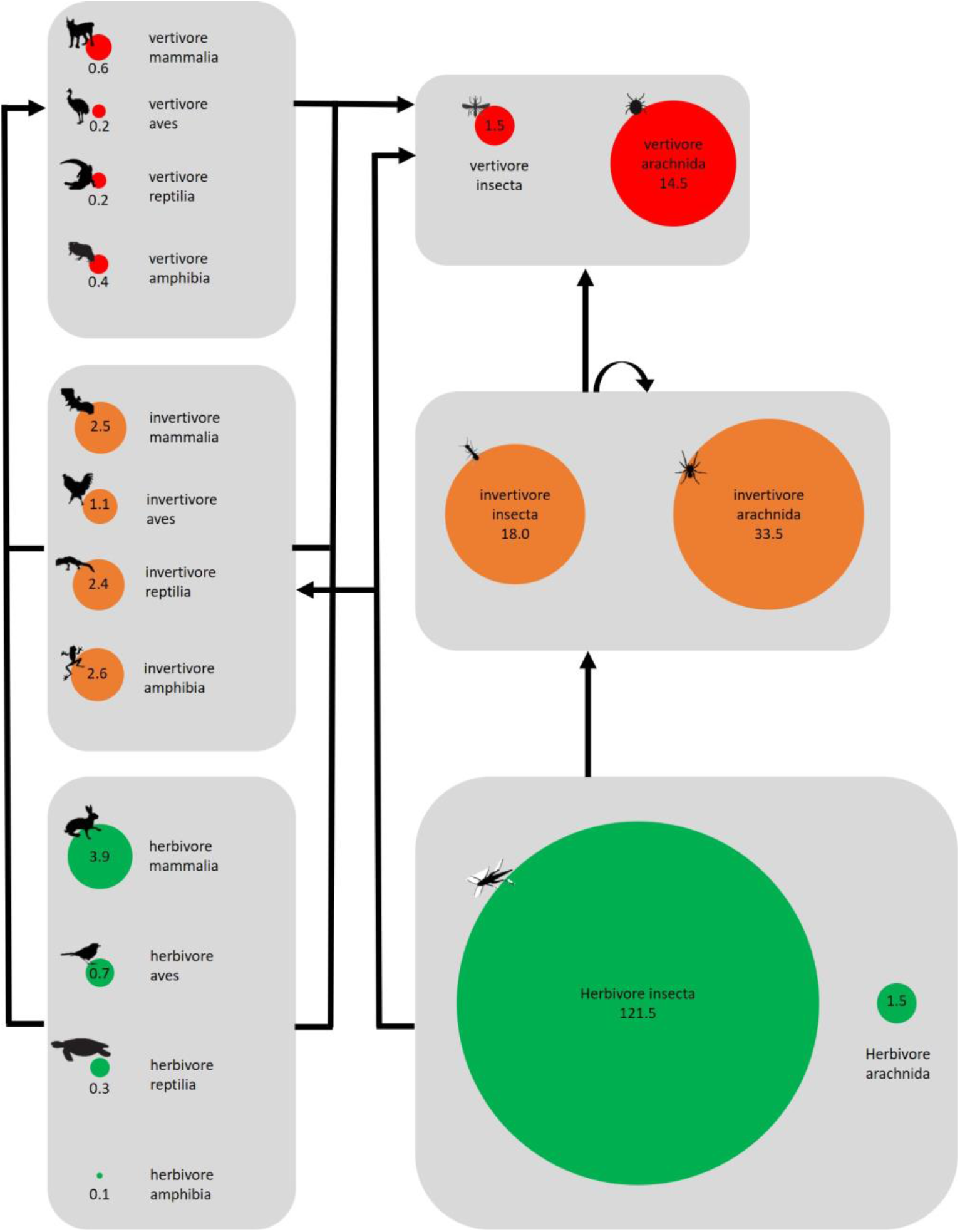
An estimated global food web for the terrestrial environment. Numbers denote the estimated global biomass (Megaton carbon) of each group, also represented by circle area. For example, among arachnida, animals with the total biomass of 33.5 Mt C are invertivores, i.e. feeding on invertebrates such as insects. Parasitic arachnida, and, to lesser extent, parasitic insects, feed on vertebrates. In this way, unexpectedly, vertebrates support a larger invertebrate biomass (16.0 Mt C) than *vice versa* (8.6 Mt C). This is so because parasitic arachnida and insects do not consume their vertebrate hosts whole; rather, they usually feed on blood of the hosts. The terrestrial top predators, vertebrates feeding on other vertebrates, make a very small fraction, 1.4 Mt C collectively. Marine mammals are included here for simplicity, although 88 species of them live exclusively in the ocean.

## Discussion

A global analysis across the animal kingdom produced a middle-heavy trophic pyramid. We confirm our hypothesis of a classic terrestrial pyramid (data not shown), while the marine pyramid was not an ‘inverted pyramid’ as hypothesized and claimed before (Buck et al. 1996, Fath and Killian 2007), but rather took the shape of a violin. Since the biomass ratio between terrestrial and marine animals is almost 1:10 (Fig. 2), the global pyramid was also middle-heavy. Still, the classic pyramid shape is true for two configurations: (1) in terms of the number of species, due to the fact that 67% of all animal species are insects, most of which are herbivores, division of all animal species among trophic levels would indeed yield a pyramid; and (2) for terrestrial animals, including terrestrial invertebrates, a pyramid of biomass holds true (not shown here). Still, we showed that animal species in most groups feed on prey, not plant. We verified our pyramid of biomass for birds, the only class with currently available body mass and abundance data, vigorously confirming our observation of only 32-36% herbivory. Interestingly, the species-weighted diet source biomass partitioning for birds yielded a much higher fraction of vertivores (45%) than our estimation (8%), further supporting our hypothesis of predation as the major dietary strategy. This might be explained by larger body sizes of birds feeding on vertebrates, rather than on invertebrates (e.g., birds of prey; Wilman et al. 2014). Other relatively large vertivore birds, like crows and seagulls, have large populations globally (Callaghan et al. 2021), also contributing to the higher vertivore biomass in birds.

The herbivore-dominated paradigm could have been biased by the prolific research on mammals on the one hand, and the lesser account of invertivores on the other hand. For example, birds of prey form just 3 out of 34 orders of birds (Brusatte et al. 2015), hiding the fact that actually most bird species (64%) feed on prey, from large aquatic birds feeding on fish, to the most delicate passerine birds feeding on invertebrates. However, perhaps the major strength of the old paradigm is its account of biomass transfer efficiency, by which each level of the pyramid must be smaller than the level below (Elton 1927, Trebilco et al. 2013). However, a major hindsight of this approach is the higher turnover of primary consumers compared to secondary consumers, and so forth (McCauley et al. 2018). On average, herbivores reproduce faster and live shorter lives than their predators, and thus can sustain a larger community of such predators. While this has already been recognized in the marine environment (Buck et al. 1996, Bar-On et al. 2018), it should explain the middle-heavy hierarchy on a global scale as well. In a global analysis as presented here, there is no longer need to define community boundaries (McCauley et al. 2018), and exogenous mechanisms of lower herbivores’ biomass become irrelevant too. Among endogenous mechanisms, higher turnover rates seem the most likely mechanism.

To the best of our knowledge, the dataset of animal diets compiled here is the largest so far. Still, while collecting data on thousands of species means a high coverage in vertebrate classes, coverage was still <1% in three invertebrate classes. However, using repeated sampling simulations, we showed that our dataset was sufficient for representing each and every class (Fig. S2). Another potential limitation of compiling a large dataset is by mixing species of unrelated ecosystems. Yet, diet partitioning across animal groups in Galapagos Islands and in Kruger National Park were similar with the global partitioning (Fig. S5). Our class-level methodology approach facilitated food source partitioning across animal classes. Still, additional uncertainties were involved in applying this approach to animal biomass. Interspecific differences in species-specific biomass and population size can affect our results; however, we assume that the role of such differences is small when averaging over hundreds and thousands of species per class.

Secondary consumers form the major link between primary consumers on the one hand, and top predators on the other hand, and hence are pivotal to almost all food-webs. Therefore, having a high biomass of secondary consumers might offer some advantage to communities and ecosystems. Considering the dual role of secondary consumers in food webs, we speculate that the middle-heavy pyramid is more resilient to change than both the classic and the inverted pyramids (Nagelkerken et al. 2020), perhaps explaining its existence on a global scale. Previous analyses and models often considered two trophic level food webs, i.e., with consumers and predators, and hence discussed either bottom-heavy or top-heavy pyramids (McCauley et al. 2018). In such cases, models showed that top-heavy pyramids were unstable in long-term simulations (although an initial stabilization phase was often identified). Here, in contrast, a three trophic levels hierarchy is intrinsic to our approach, including herbivores, invertivores, and vertivores. Doing so consistently, opens up the third possibility of middle-heavy hierarchy. Here we argue that such a pyramid can be stable and prevalent.

The potentially higher resilience of the middle-heavy pyramid could be important at the current age of the Anthropocene. We assume that the number of species extinct due to human activity, although alarmingly significant, is still small relative to the number of extant species (Ceballos et al. 2010). This considered, at the species level, our middle-heavy pyramid should represent the native condition. We further confirmed the dominance of prey over plant by observations from two native ecosystems (Figs S3, S4). Yet at the biomass level, the biomass estimates used here are based on current censuses, and hence already reflect animal populations that have been altered, and mostly decreased, by humans. The native, pre-Anthropocene, biomass of each animal group is virtually unknown. Notably, defaunation is not homogeneous across animal groups (McCauley et al. 2015): for example, humans feed on specific wild animals and not on others, and even at the group level, mammals, birds, and fish are traditional human prey, whereas reptiles, amphibians, arachnida and cnidaria, much less so. Anthropogenic effects are also biasing natural food webs, in ways that typically affect higher trophic levels more than lower levels (Sandin et al. 2008, McCauley et al. 2015). It was hence speculated that pre-Anthropocene ecosystems were more top-heavy than today (Sandin et al. 2008). In addition, direct consumption (hunting, fishing) is probably the least among anthropogenic means of defaunation, like habitat loss and climate change. Human-induced animal invasions have been shown to reduce predator biomass in some ecosystems (McCauley et al. 2012), while increasing predator biomass in other instances, e.g. the introduction of foxes in subarctic islands (Maron et al. 2006). Evidently, effects on specific species propagate into effects on additional species through the food web. These secondary effects are not linear, nor homogeneous: a decrease in the population of one species will not only decrease the entire food web around it, but rather change the whole organization of the trophic pyramid (Nagelkerken et al. 2020). Overall, it is certainly possible that the middle-heavy pyramid found here, both globally and locally, holds true not only at the species level but also at the biomass level. Since it is safe to assume that humanity has put more pressure on higher than lower trophic levels, a native global fauna would have probably been richer in vertivores and invertivores than today, and not the opposite.

Why do most insects and mammals, unlike most animals, feed on plants? Evidence show that predation, and not herbivory, is the primitive habit: the first ancestor of all animals was a carnivore eating other heterotrophs, and the ability to feed on plants was a later habit, not without added value (Román-Palacios et al. 2019). Among extant animals, mammals are notably larger than most animals in the other groups, forming the planet’s megafauna (Asner et al. 2016). Body mass varies across mammals due to energetic constraints: larger animals can behaviorally obtain and ingest proportionally more food than smaller animals (Gittleman 1985). Larger animals have lower metabolic rates, and hence can feed on low-energy food which is slow to release energy in digestion, whereas small animals need to select food with high energy and nutrition, like prey. Small animals are also limited in the size of prey they can feed on (e.g., insects can rarely feed on vertebrates, unless parasitic; Fig. S7). In mammals, across the order Carnivora, diet type is the major explanatory factor for body mass, where herbivores (in the order carnivora) are the largest, followed by piscivores, carnivores, and the smaller insectivores (Gittleman 1985). In addition, bite force (normalized for body mass) is highest for both carnivores preying on large prey and herbivores consuming tough fibrous plant material, and lowest in insectivores (Christiansen and Wroe 2007). To test if herbivory is related to higher body mass and lower metabolic rate, we collected data on food sources of dinosaurs, the previous megafauna group on Earth. Indeed, plants contributed 62% of dinosaur diet (Fig. S8; anecdotally still lower than the 76% of modern human diet). Insects, too, are claimed to have once formed a megafauna group, with evidence of giant insects ∼300 million years ago. However, insect herbivory could be more readily explained by other traits: (1) the small size and low speed of many insects limit their options as predators (although Hymenopterans are major carnivores and parasitoids) (Schoenly 1990; Fig. S7); (2) insects are ultimate plant feeders in terms of speed of consumption and efficiency; and (3) insect metamorphosis means that most feeding is completed in the juvenile stage. We note, in passing, that this research takes a snapshot, as it were, of partitioning of animal diets, at the present time. Animal groups have evolved, appearing and disappearing on the stage of the Earth, throughout millions of years. One curious example could be the relationship between tarantula spiders and birds. The former seem to have appeared in the fossil record in the Triassic, before the appearance of birds. When birds did appear, in the Late Jurassic, spiders may have been afforded an avian addition to their diet. Later on, some bird species may have evolved to feed on tarantulas.

Finally, a major implication from our study is that, for most animals (by biomass), plants are not a direct food source. Instead, animals in most groups rely on invertebrates for their food, and, in the case of terrestrial animals, on insects (Fig. 4). Indeed, insects have already been recognized as important in the evolution of food web complexity (Schoenly 1990). To predators, insects offer a highly nutritious food, which is higher in abundance than any other prey; easier to catch, kill, and chew (Gittleman 1985, Christiansen and Wroe 2007). Insects are a source of energy, protein, fat, minerals and vitamins, with the energy content being on a par with vertebrate meat sources (Finke 2002, Dobermann et al. 2017). Studies with juvenile rats have demonstrated that crickets (*Acheta domesticus*) offer a superior source of protein when compared to a plant source (soy protein) (Belluco et al. 2013). Among terrestrial vertebrates, early insectivory is supported by arthropod remains in the oral cavities of fossil reptiles (Modesto et al. 2009). Analysis of jaw shape of fossils shows that most early (Mesozoic) mammals were insectivores, and a few taxa were vertivores (Morales-García et al. 2021). The key role of insects in the food web is demonstrated by parallel declines in abundance of insects and insectivorous birds in Denmark over 22 years, related with pesticide use (Møller 2019). Essentially, a world without insects would potentially mark the end of complex life on Earth.

## Acknowledgments

The authors wish to thank the many experts who provided advice on the diverse topics covered in the analysis: Yinon Bar-on, Ron Milo, Gitai Yahel, Uri Roll, Shai Meiri, Michal Segoli, Moshe Coll, Sharoni Shafir, Eshel Ophir, Eyal Krieger, Amatzia Genin, Bracha Farstey, Kristin Palmqvist, Boaz Yuval, David Lerner, Ido Rog, Ella Pozner, Adva Shemi, Nicolas Bailly, Daniel Pauly, Roslyn Cameron, Izak Smit, Guin Zambatis, Roi Holzman, and Uri Alon. Animal silhouette icons for figures were taken from vectorstock.com

## Author contributions

TK designed the study and UM collected the data. SLL performed the statistical analysis. TK performed the data analysis and wrote the manuscript.

## Data and materials availability

The full dataset will be made available on Figshare.com following acceptance of the paper.

## Supplementary Information

### Supplementary Figures

**Fig. S1.**
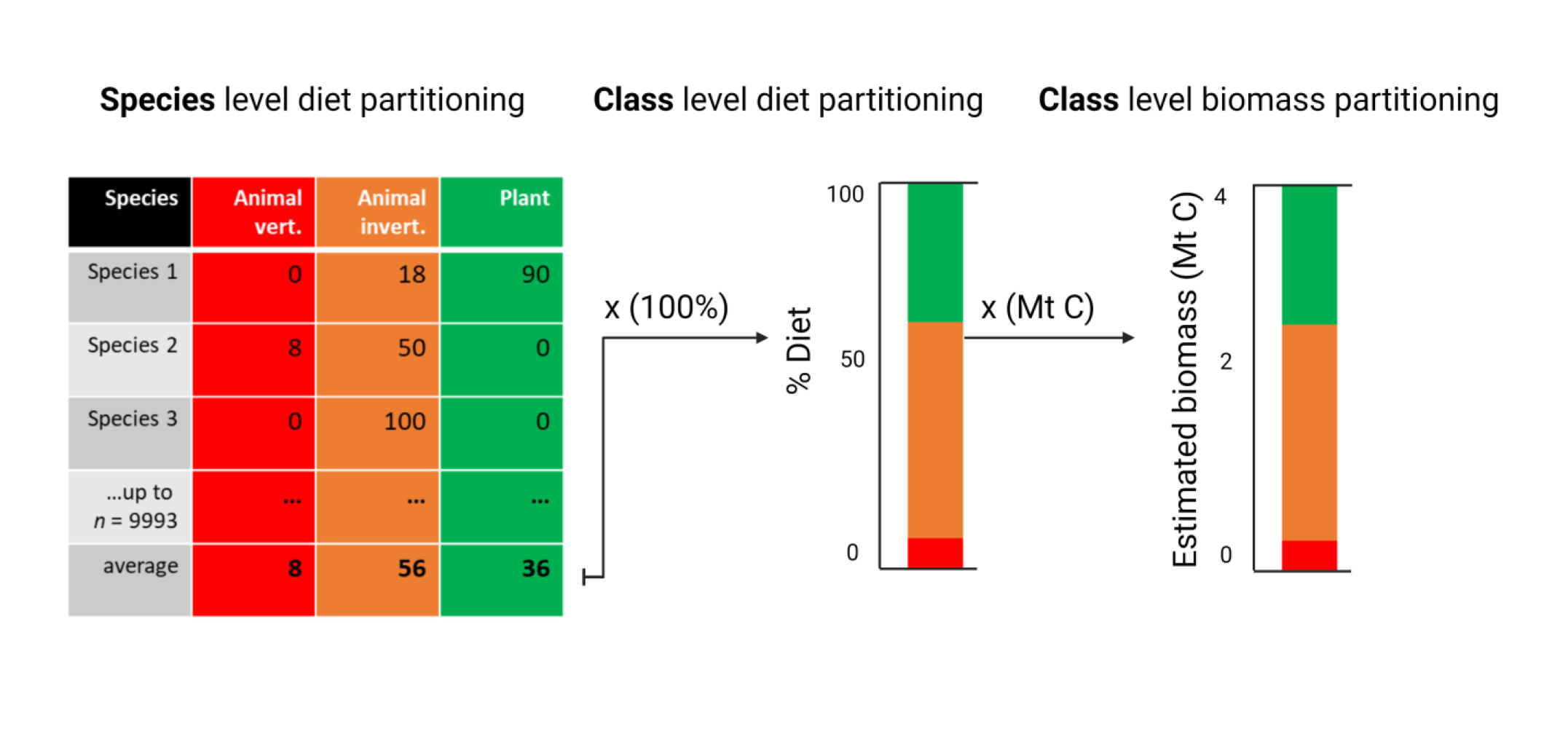
A class-level methodology approach facilitates food source partitioning across animal classes. In this study we used species-level diet partitioning as surrogate for species-specific biomass and population size, which are not available (Table 1). Species-level diet partitioning was used for a class-level diet partitioning by equally weighted averaging. Next, we multiplied the class-level partitioning by a class-level biomass estimate, to produce a biomass estimate for the role of each food source in feeding each of the animal groups.

**Fig. S2.**
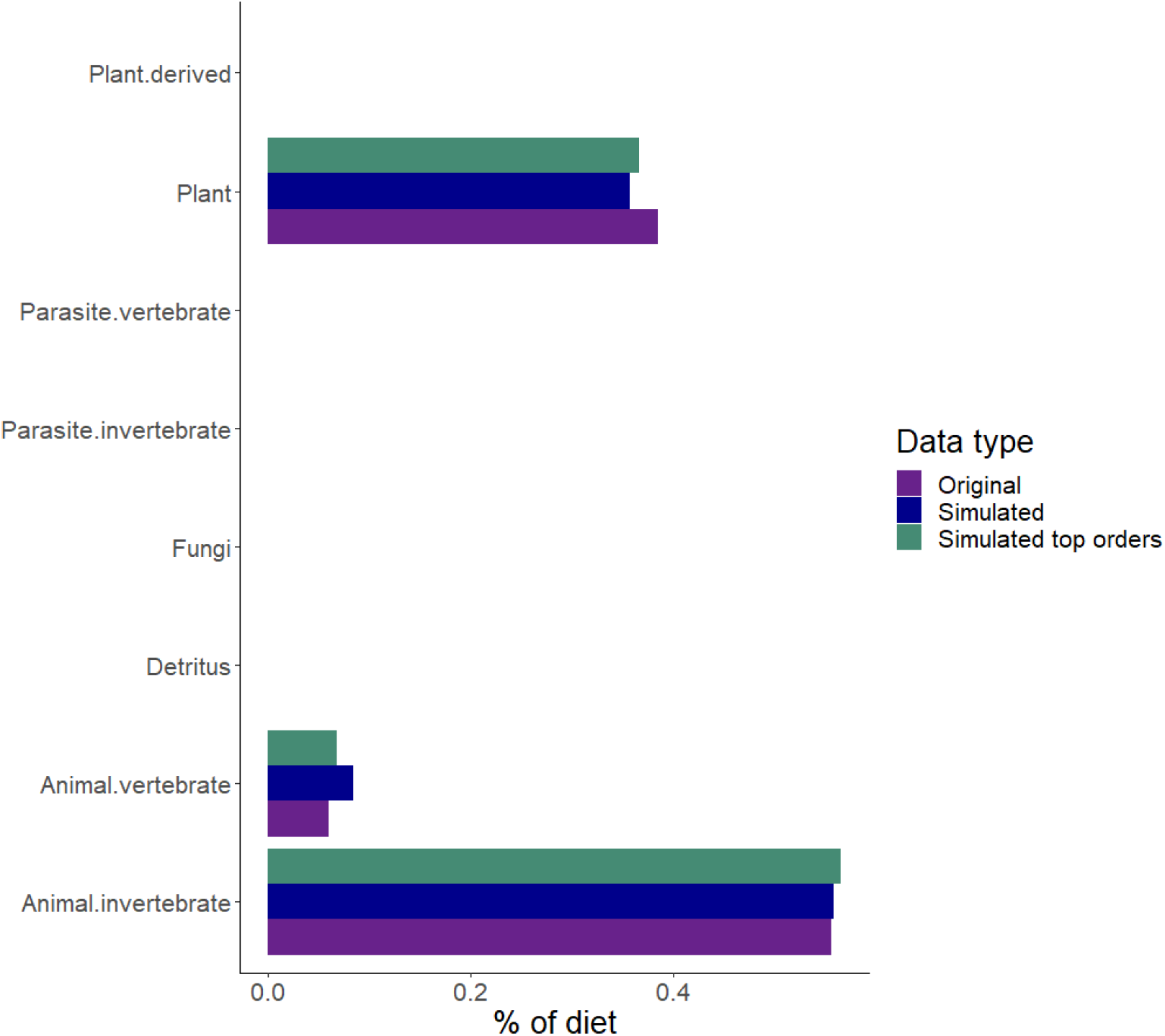
The low sampling rate among insects is sufficient to represent their diet partitioning, as tested and confirmed among bird species. Diet partitioning in the original complete dataset of birds (purple, *n*=9994) and for the simulated data (mean of 10,000 repeats, *n*=30) including all of birds’ orders (i.e., a random sampling; blue) or only the top birds’ orders, collectively containing 93% of all bird species (green, *n*=30).

**Fig. S3.**
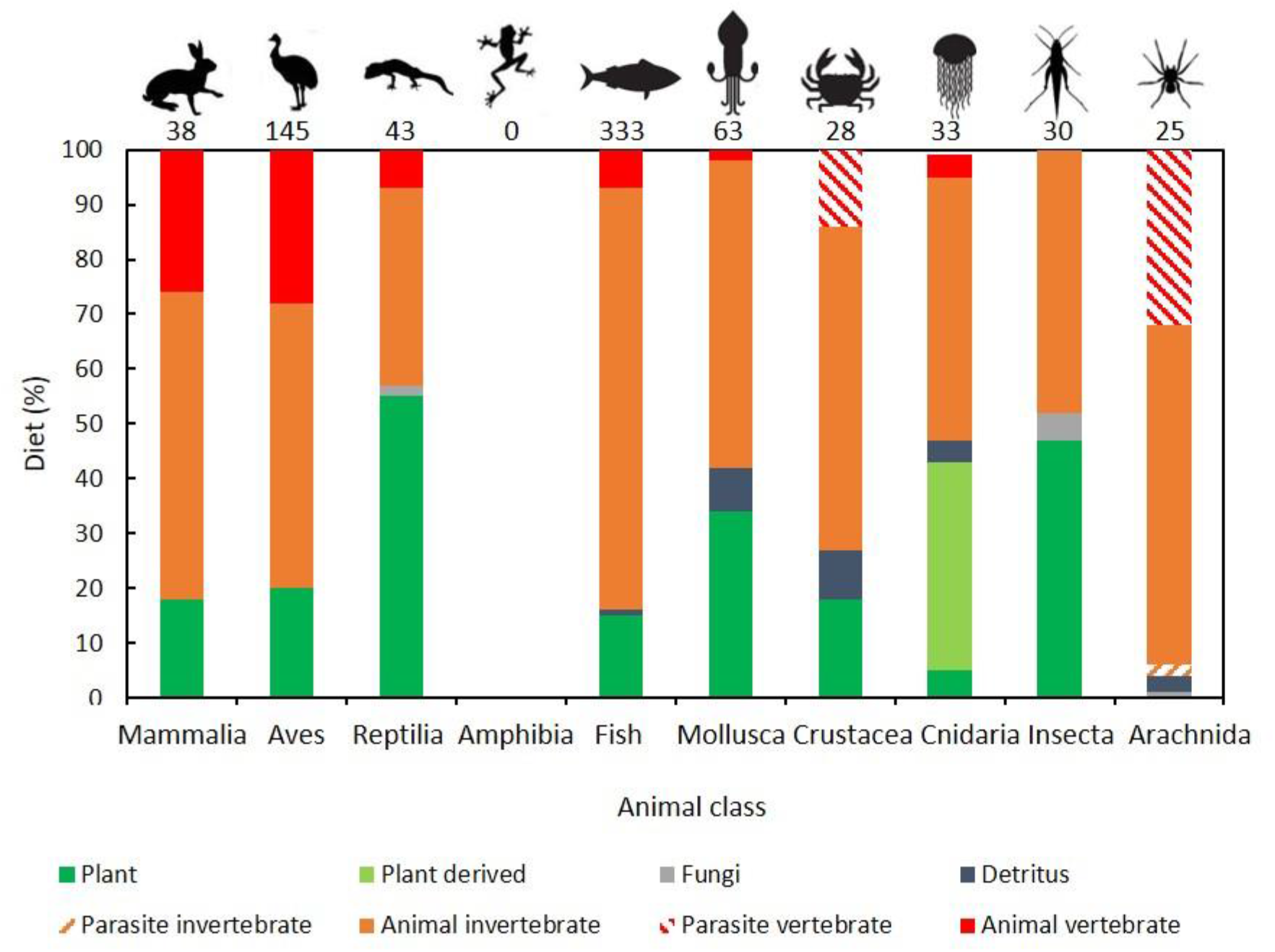
Galapagos Islands species in most animal groups feed on prey, not plant. Partitioning of food sources among nine major animal classes (no native amphibian species in Galapagos). Numbers on top indicate the number of sampled species in each class/ subphylum/ phylum. Fish include all Chondrichthyes and Osteichthyes species.

**Fig. S4.**
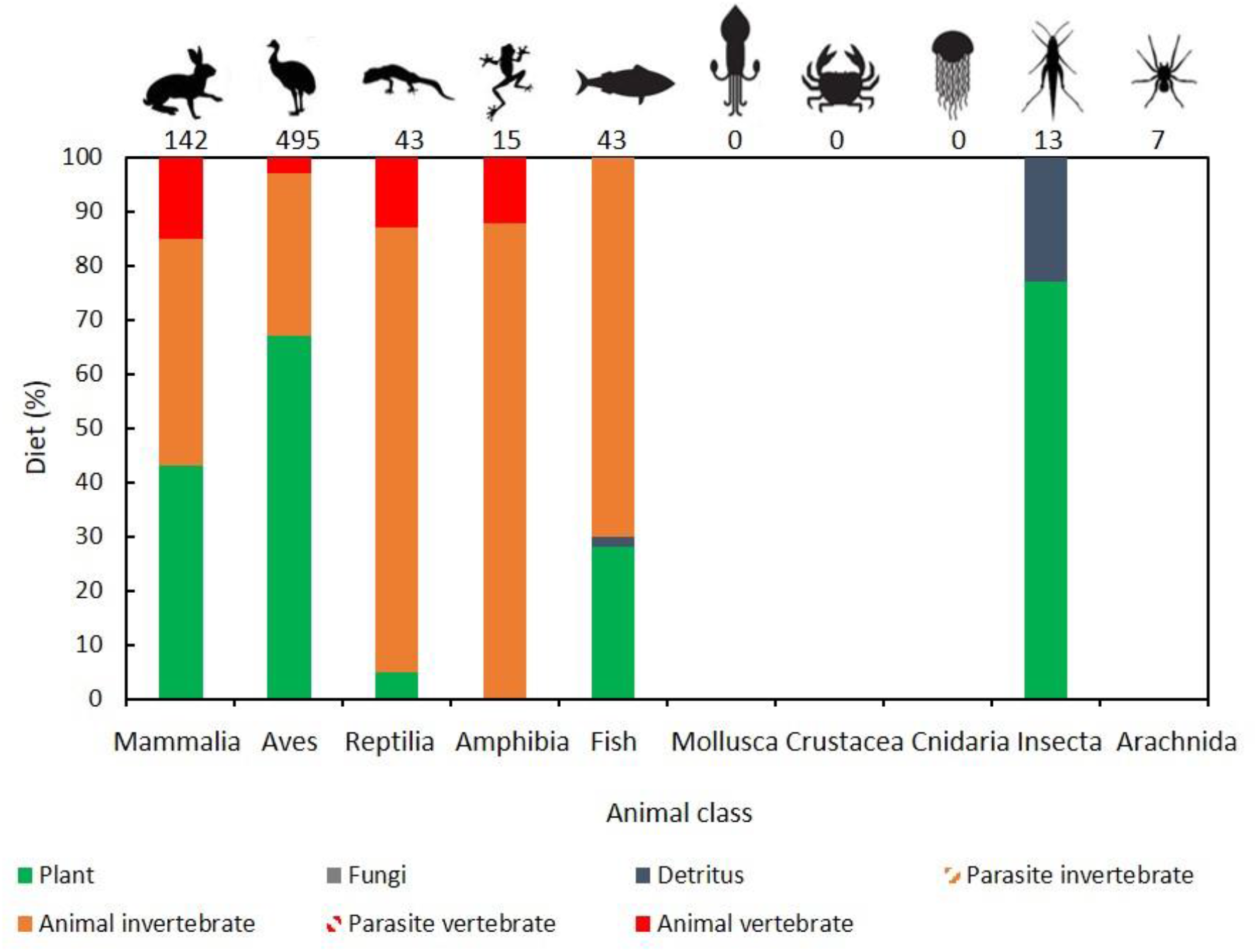
Kruger National Park species in most animal groups feed on prey, not plant. Partitioning of food sources among six major animal classes (native marine invertebrate classes had too low number of species in Kruger; diet data missing for arachnida species). Numbers on top indicate the number of sampled species in each class/ subphylum/ phylum. Fish include all Chondrichthyes and Osteichthyes species.

**Fig. S5.**
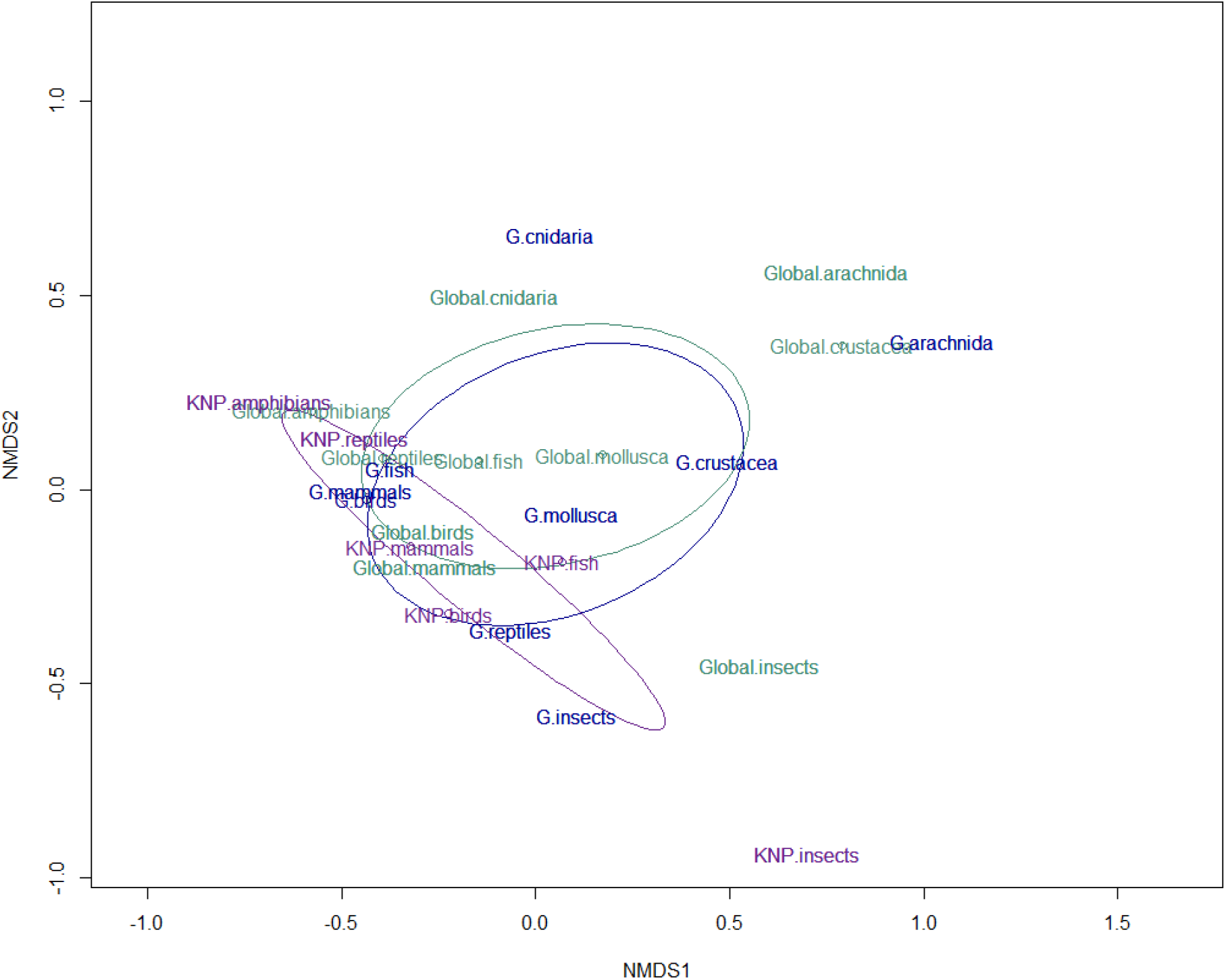
Diet partitioning across animal groups in Galapagos Islands and in Kruger National Park are similar with the global partitioning. Non-metric multidimensional scaling (NMDS) based on Bray-Curtis dissimilarities among the diet partitioning of different animal groups in the various datasets examined: the global dataset (green), the Galapagos dataset (blue) and the Kruger dataset (purple).

**Fig. S6.**
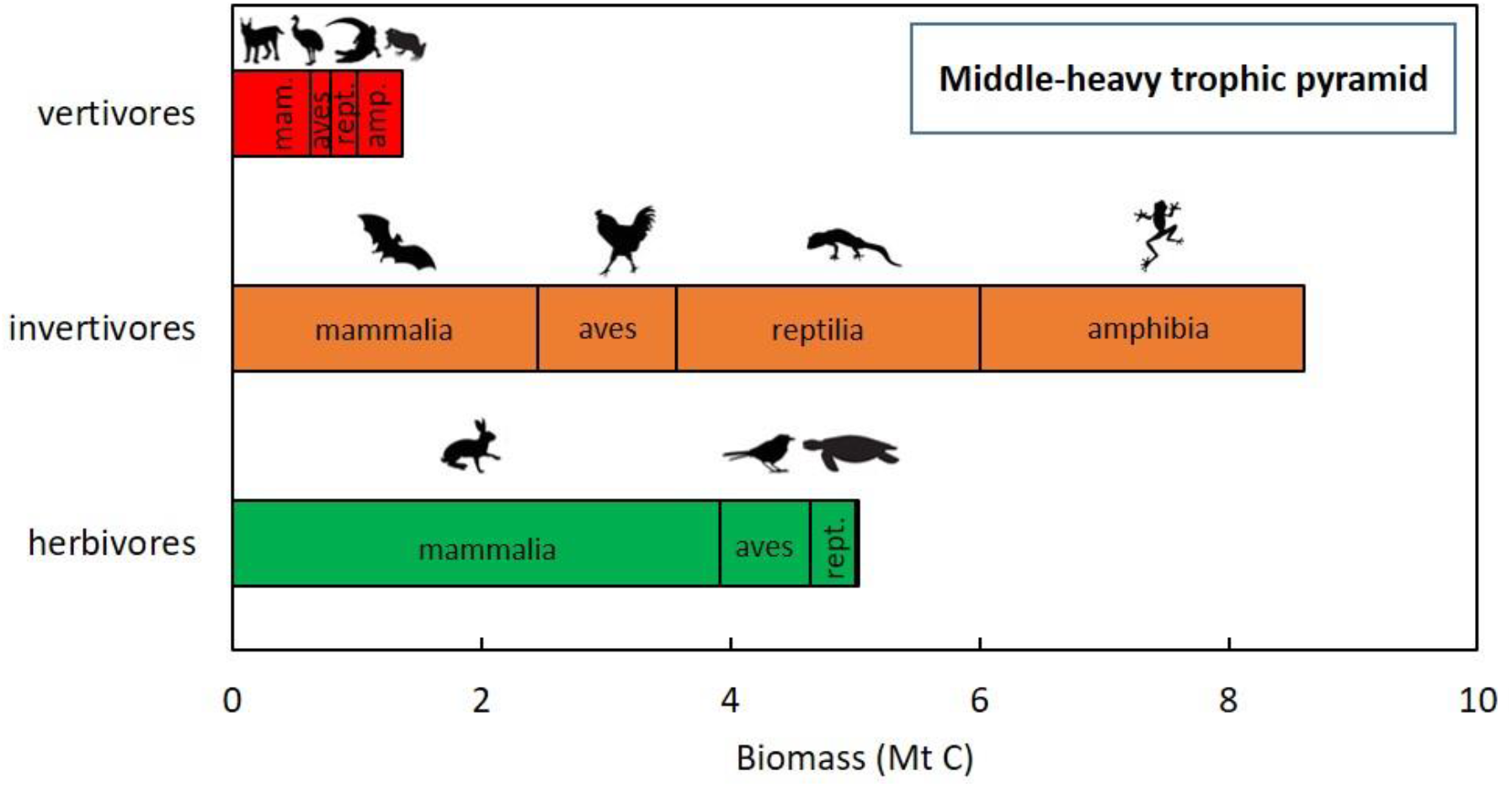
The trophic pyramid of terrestrial vertebrates is middle-heavy: invertivores (secondary consumers) are more abundant than herbivores (primary consumers). Estimated biomass (Megaton carbon) of herbivores, invertivores, and vertivores of each of the four non-fish vertebrate groups (mam., mammalia; rept., reptilia; amp, amphibia).

**Fig. S7.**
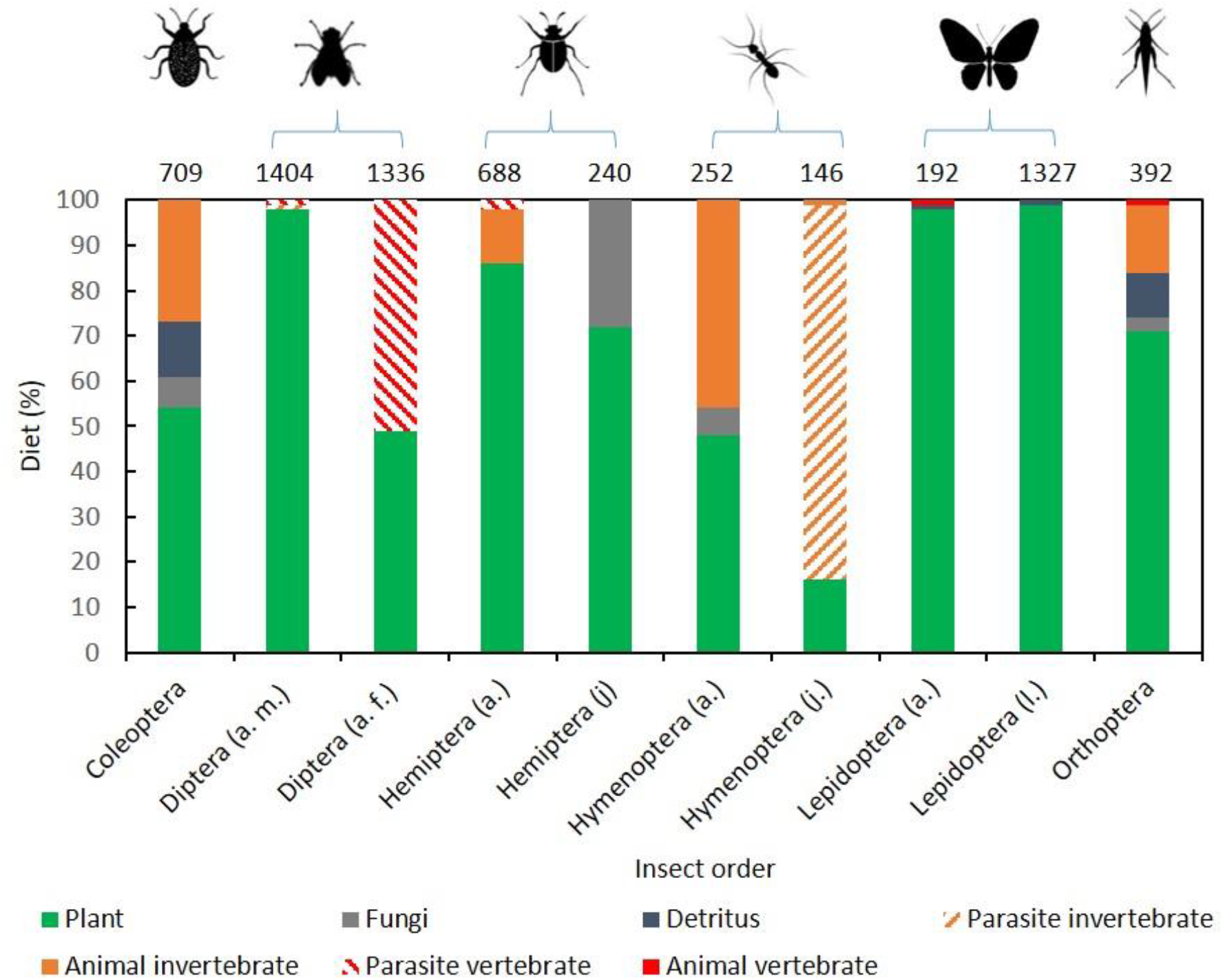
Species in most insect orders feed on plants. Partitioning of food sources among six major insect orders. a, adults; j, juveniles; l, larva; f, female; m, male. Numbers on top indicate the number of sampled species in each order.

**Fig. S8.**
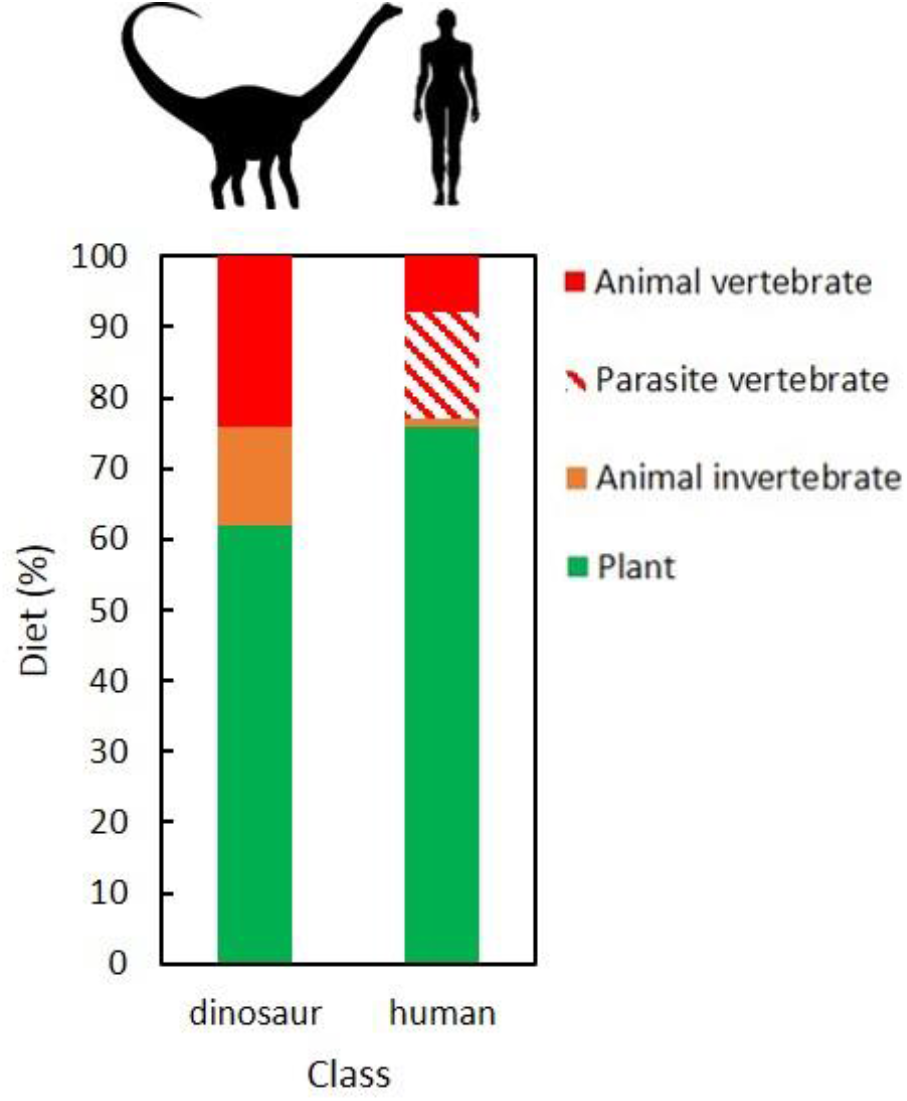
Dinosaur species and humans feed mostly on plants. Partitioning of food sources among 102 dinosaur species and among modern humans (a global average).

